# Human GBP1 promotes pathogen vacuole rupture and inflammasome activation during *Legionella pneumophila* infection

**DOI:** 10.1101/2020.05.27.120477

**Authors:** Antonia R. Bass, Sunny Shin

## Abstract

The inflammasome is an essential component of host defense against intracellular bacterial pathogens, such as *Legionella pneumophila*, the causative agent of the severe pneumonia Legionnaires’ disease. Inflammasome activation leads to recruitment and activation of caspases, which promote IL-1 family cytokine release and pyroptosis. In mice, interferon (IFN) signaling promotes inflammasome responses against *L. pneumophila*, in part through the functions of a family of IFN-inducible GTPases known as guanylate binding proteins (GBPs) (1). Within murine macrophages, IFN signaling promotes rupture of the *L. pneumophila*-containing vacuole (LCV), whereas GBPs are dispensable for vacuole rupture. Instead, GBPs facilitate the lysis of cytosol-exposed *L. pneumophila*. In contrast to mouse GBPs, the functions of human GBPs in inflammasome responses to *L. pneumophila* are poorly understood. Here, we show that IFN-γ promotes caspase-1, caspase-4, and caspase-5 inflammasome activation during *L. pneumophila* infection and upregulates GBP expression in primary human macrophages. We find that human GBP1 is important for maximal IFN-γ-driven inflammasome responses to *L. pneumophila*. Furthermore, IFN-γ signaling promotes the rupture of LCVs. Intriguingly, in contrast to murine GBPs, human GBP1 targets the LCV in a T4SS-dependent manner and promotes vacuolar lysis, resulting in increased bacterial access to the host cell cytosol. Our findings show a key role for human GBP1 in targeting and disrupting pathogen-containing vacuoles and reveal mechanistic differences in how mouse and human GBPs promote inflammasome responses to *L. pneumophila*.

## Introduction

The innate immune response to bacterial pathogens is essential for mediating host defense and bacterial clearance. This response is initiated through the recognition of conserved microbial components known as pathogen-associated molecular patterns (PAMPs) by host pattern recognition receptors (PRRs) (2, 3). In particular for intracellular bacteria, a subset of cytoplasmic PRRs that detect bacterial components contaminating the host cell cytosol and other activities associated with invading pathogens has been implicated in host defense. Upon activation, host sensors such as the nucleotide-binding oligomerization domain-like receptors (NLRs) mediate the formation of a multimeric protein complex termed the inflammasome. Inflammasome activation triggers a cascade of immune responses that culminate in the release of IL-1 family cytokines and an inflammatory form of cell death termed pyroptosis. This response alerts the body of the infection and recruits other innate immune cells to the site of infection, thereby promoting bacterial control and clearance.

The two major inflammasomes that have been described are the canonical and noncanonical inflammasomes. In response to a diverse range of ligands, canonical inflammasomes recruit and activate the cysteine protease caspase-1 to promote the processing and secretion of the proinflammatory cytokines IL-1β and IL-18 (4, 5). Additionally, an alternative caspase-1-independent inflammasome termed the noncanonical inflammasome mediates inflammatory responses to gram-negative bacteria (6–12). The noncanonical inflammasome is formed by caspase-11 in mice and two orthologs in humans, caspase-4 and caspase-5; these caspases are activated upon binding bacterial lipopolysaccharide (LPS), a potent PAMP and major outer membrane lipid component of gram-negative bacteria (13–15). Following their activation, these inflammatory caspases cleave the substrate gasdermin-D (GSDMD). Upon cleavage, the GSDMD N-terminal fragment translocates to the plasma membrane and oligomerizes to form a pore, leading to pyroptosis (16, 17). Death of the infected cell eliminates the replicative niche for intracellular pathogens, and results in uptake of the bacteria within pore-induced intracellular traps (PITs) by neutrophils and subsequent clearance of the bacteria by efferocytosis *in vivo* (18).

Inflammasome responses are potentiated by priming signals recognized by plasma membrane receptors that upregulate the production of inflammatory cytokines and inflammasome components. During an infection, toll-like receptors play a major role in promoting the expression of innate immune genes. Additionally, type I and type II IFNs produced during infection promote inflammasome responses in mice. A subfamily of IFN-upregulated GTPases called GBPs are particularly important in promoting inflammasome responses to gram-negative bacteria in mice (19–28). Mouse GBPs can localize to pathogen-containing vacuoles (29). However, the precise steps regulated by GBPs in promoting inflammasome activation are unclear. Earlier studies with *Salmonella* Typhimurium indicated that GBPs promote rupture of pathogen-containing vacuoles (PCVs), whereas later studies with *Francisella novicida* and *L. pneumophila* indicate that GBPs function downstream of PCV rupture and facilitate bacteriolysis, resulting in cytosolic release of bacterial components that subsequently trigger inflammasome activation (20–22, 30, 31). Mouse GBPs can also promote inflammasome responses in the absence of binding to the PCV, as is the case with the vacuolar pathogen *Chlamydia muridarum* (32). It is still unclear how mouse GBPs mediate these various functions, although one study showed that GBPs recruit the immunity-related GTPase (IRG) IRGB10 to mediate bacteriolysis (30).

While studies in mice have linked the functions of IFN signaling and GBPs to inflammasome activation, the degree to which the function of murine GBPs mirror their human counterparts is unknown, as the significant differences in immune genes between mice and humans, including in the GBP superfamily, could translate into differences in immune mechanisms. Notably, mice have 11 GBPs, whereas humans only have seven GBPs (33). The functions of human GBPs in host defense against gram-negative bacteria, particularly whether human GBPs play a role in PCV rupture or bacteriolysis, is unclear. Human GBP1 binds to the outer membrane of the cytosolic pathogen *Shigella flexneri* and further recruits additional GBPs, specifically GBP2, 3, 4, and 6, to inhibit the actin-based motility of *S. flexneri* (34, 35). Additionally, human GBP1 and GBP5 promote inflammasome responses to the vacuolar pathogen *S*. Typhimurium, while human GBP2 promotes inflammasome responses to the cytosolic pathogen *F. novicida* (27, 36, 37). These findings indicate that different human GBPs function in a bacterium-specific manner.

Here, we sought to define the role of IFN-γ signaling and human GBPs in human inflammasome responses to the vacuolar pathogen *L. pneumophila*. *L. pneumophila* is a gram-negative intracellular bacterial pathogen that infects alveolar macrophages and is the causative agent of the severe pneumonia known as Legionnaires’ Disease (38). Upon uptake, *L. pneumophila* resides within a *L. pneumophila*-containing vacuole (LCV) and relies on the Dot/Icm type IV secretion system (T4SS) to survive within the LCV (39–44). The T4SS injects over 300 effector proteins, many of which enable *L. pneumophila* to evade the endolysosomal pathway and modify its LCV into an ER-derived replicative compartment (45–50). Despite being essential for *L. pneumophila* virulence, T4SS activity triggers robust canonical and noncanonical inflammasome activation in human macrophages (51). The role of IFN signaling and GBPs in promoting human inflammasome responses to *L. pneumophila* is unknown.

In this study, we found that IFN-γ promotes inflammasome responses to *L. pneumophila* in a T4SS-dependent manner in both immortalized and primary human macrophages. We further determined that human GBP1 was essential for maximal inflammasome activation and that IFN-γ-primed macrophages had a significant increase in GBP1 and GBP2 localization to the LCV compared to unprimed macrophages. GBP1 and GBP2 were recruited to *L. pneumophila* in a T4SS-dependent manner, indicating that human GBPs detect pathogen-containing vacuoles containing virulence-associated bacterial secretion systems. Additionally, IFN-γ treatment led to the increased rupture of LCVs and exposure of *L. pneumophila* to the host cell cytosol, in part through a mechanism involving GBP1. Overall, our findings indicate that IFN-γ-dependent human GBP1 responses promote rupture of the LCV, facilitating bacterial detection in the cytosol to enhance inflammasome activation. Furthermore, as human GBP1 facilitates LCV rupture, in contrast to mouse GBPs, which are dispensable for LCV rupture, our findings suggest that mouse and human GBPs have evolved distinct functions.

## Results

### IFN-γ promotes inflammasome activation in human macrophages during *L. pneumophila* infection

IFN-γ promotes human inflammasome responses to the cytosolic pathogen *F. novicida* (37). However, whether IFN-γ upregulates inflammasome responses to a vacuolar pathogen in human macrophages is poorly understood; therefore, we sought to test this with *L. pneumophila*. To determine whether IFN signaling increases inflammasome activation in response to *L. pneumophila*, we primed macrophages with IFN-γ prior to infection with *L. pneumophila*. *L. pneumophila* requires a T4SS to translocate bacterial products into the host cell cytosol; therefore, we also investigated whether IFN-γ-mediated inflammasome responses to *L. pneumophila* are dependent on its T4SS. Unprimed or IFN-γ-primed phorbol 12-myristate 13-acetate (PMA)-differentiated THP-1 macrophages were infected with a *L. pneumophila dotA* mutant lacking a functional T4SS (T4SS-) or a T4SS-sufficient (T4SS+) strain lacking flagellin (Δ*flaA*) in order to focus on NAIP-independent inflammasome responses. Unprimed THP-1 cells infected with T4SS-*Lp* or mock infected exhibited little to no cell death, whereas cells infected with T4SS+ *Lp* underwent increased cell death and IL-1 family cytokine release (Fig. 1A and B), consistent with previous findings showing that *L. pneumophila* induces T4SS-dependent inflammasome responses in THP-1 cells (51). THP-1 macrophages that were primed with IFN-γ and infected with T4SS+ *Lp* had a significant increase in cell death compared to unprimed macrophages (Fig. 1A and S1A). IFN-γ-primed macrophages infected with T4SS+ *Lp* also had significantly elevated levels of IL-1β and IL-18 secretion compared to unprimed macrophages (Fig. 1B). Interestingly, we noticed significantly increased secretion of IL-1β and IL-18 levels in T4SS-*Lp*-infected THP-1 cells primed with IFN-γ compared to unprimed cells, although at lower levels than those observed in T4SS+ *Lp*-infected THP-1 cells primed with IFN-γ. Furthermore, we observed processing of IL-1β into its mature p17 form in the supernatant of both T4SS-and T4SS+ *Lp*-infected THP-1 cells primed with IFN-γ (Fig. 1C). These data indicate that IFN-γ priming promotes inflammasome responses to both T4SS- and T4SS+ *L. pneumophila* in THP-1 cells, although maximal inflammasome activation occurs in primed cells infected with bacteria that harbor a functional T4SS.

**Fig. 1.**
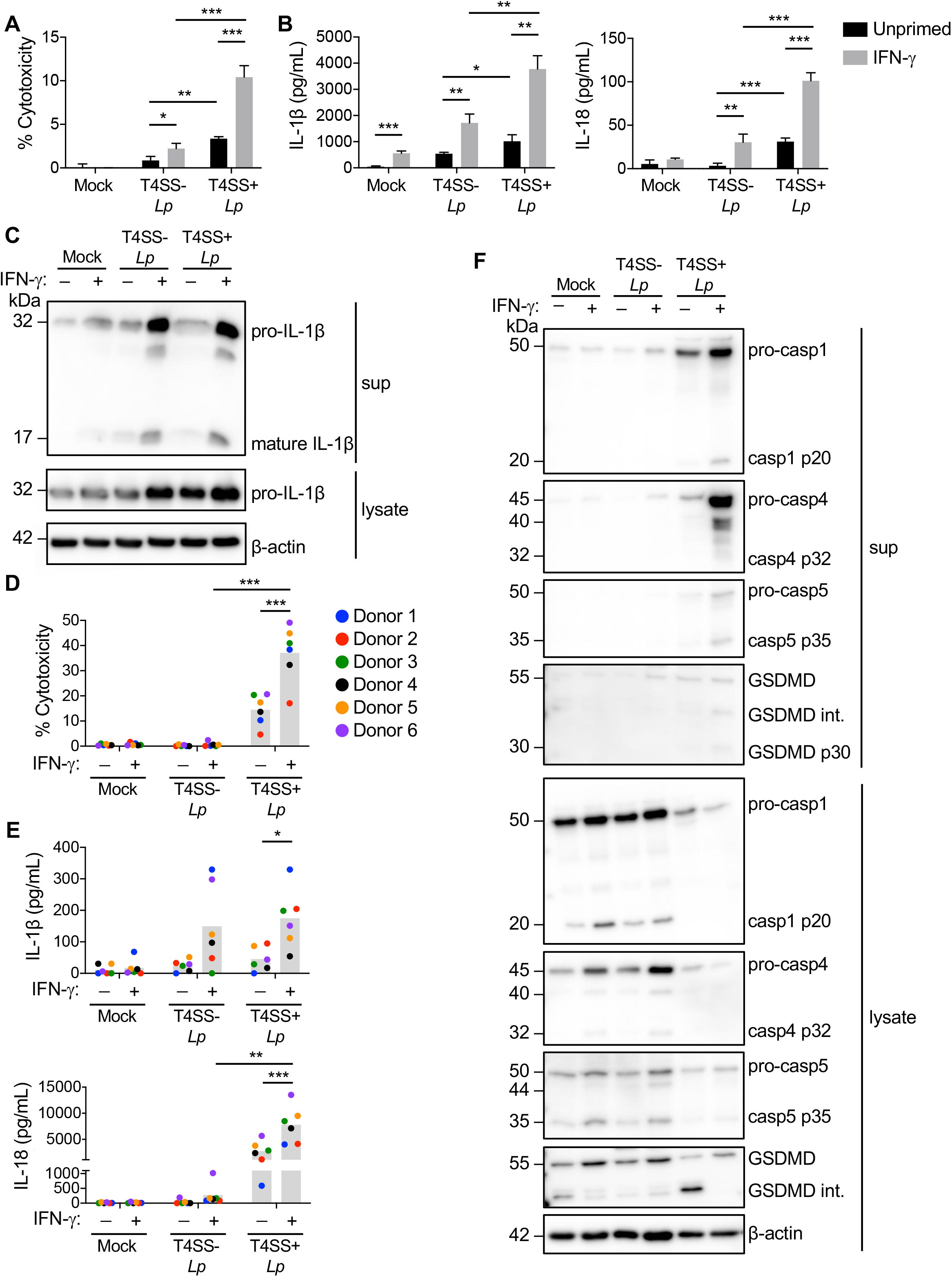
IFN-γ promotes inflammasome activation in response to *L. pneumophila* in human macrophages. Phorbol 12-myristate 13-acetate (PMA)-differentiated THP-1 cells (A, B, C) or primary human monocyte-derived macrophages (hMDMs) (D, E, F) were either left unprimed or primed with IFN-γ (100 U/ml) overnight and infected with T4SS-*Lp*, T4SS+ *Lp*, or mock-infected with PBS for two or four hours, respectively. (A and D) Cell death was measured using lactate dehydrogenase release assay and normalized to mock-infected cells. (B and E) IL-1β and IL-18 levels in the supernatant were measured by ELISA. (C, F) Immunoblot analysis was conducted on supernatants (sup) and lysates from THP-1 cells (C) or hMDMs (F) for full-length IL-1β (pro-IL-1β), cleaved IL-1β (mature IL-1β), full length caspase-1 (pro-casp1), cleaved casp1 (casp1 p20), pro-casp4, cleaved caspase-4 (casp4 p32), pro-casp5, casp5 p35, full-length Gasdermin-D (GSDMD), intermediate and cleaved GSDMD (GSDMD int. and GSDMD p30), and β-actin. Western blots are representative of three independent experiments. (A and B) Shown are the results representative of three independent experiments. *P< 0.05, **P< 0.01, and ***P<0.001 by unpaired t-test. (D and E) Shown are the pooled results of six independent experiments using hMDMs from different healthy human donors. Each data point represents the mean of triplicate infected wells from an individual donor. *P< 0.05, **P<0.01, and ***P< 0.001 by paired t-test.

We next asked whether IFN-γ also enhances inflammasome responses to *L. pneumophila* in primary human monocyte-derived macrophages (hMDMs) derived from healthy human donors. IFN-γ-primed hMDMs infected with T4SS+ *Lp* also exhibited significantly increased levels of cell death (Fig. 1D), as well as IL-1β and IL-18 release (Fig. 1E), compared to unprimed or IFN-γ-primed hMDMs that were uninfected or infected with T4SS-*Lp*. Overall, our data indicate that IFN-γ promotes inflammasome responses and IL-1 family cytokine release in response to *L. pneumophila* infection in both PMA-differentiated THP-1 cells and primary hMDMs.

### Caspase-1 and additional caspases promote inflammasome activation in response to *L. pneumophila*

We next investigated which caspases are involved in promoting inflammasome activation in response to *L. pneumophila* following IFN-γ priming. *L. pneumophila* activates the human noncanonical caspase-4 inflammasome in unprimed macrophages (51), but whether IFN-γ priming affects canonical or noncanonical inflammasome activation in *L. pneumophila*-infected human macrophages has not been studied. We observed caspase-1 processing into its mature p20 form in T4SS+ *Lp*-infected hMDMs primed with IFN-γ (Fig. 1F). Both caspase-4 and caspase-5 were upregulated at the RNA and protein level following IFN-γ priming of THP-1 cells and hMDMs (Fig. 1F and S1B-D). Additionally, we observed release of full-length and processed forms of caspase-4 and caspase-5, as well as GSDMD processing and release, into the supernatants of IFN-γ-primed hMDMs infected with T4SS+ *Lp* (Fig. 1F). Together, these data demonstrate that caspase-1, −4, and −5 are processed into their mature forms upon IFN-γ priming and infection with T4SS+ *Lp* (Fig. 1F).

We next tested whether caspase activity is required for inflammasome responses. Cell death was significantly decreased in either unprimed or IFN-γ-primed hMDMs that were treated with the pan-caspase inhibitor ZVAD prior to infection with T4SS+ *Lp*, compared to the levels of cell death observed in vehicle control-treated cells (Fig. 2A). IL-1β and IL-18 secretion was also significantly decreased in IFN-γ-primed hMDMs treated with ZVAD, compared to DMSO-treated hMDMs (Fig. 2B and C). In addition, IL-18 secretion was significantly decreased in unprimed hMDMs treated with ZVAD. Importantly, treatment with the caspase-1-specific inhibitor YVAD significantly reduced cell death and IL-1β and IL-18 secretion in IFN-γ-primed hMDMs, compared to DMSO-treated hMDMs. Similarly to ZVAD treated cells, YVAD also significantly reduced cell death and IL-18 release in unprimed hMDMs. Interestingly, we observed lower amounts of cell death and IL-1 family cytokine release in hMDMs treated with the broader-spectrum inhibitor ZVAD compared to treatment with the caspase-1 selective inhibitor. These data indicate that caspase-1 and likely additional caspases are involved in promoting inflammasome responses to *L. pneumophila*. As both caspase-4 and caspase-5 are processed in IFN-γ-primed T4SS+ *Lp*-infected hMDMs (Fig. 1F), these noncanonical inflammatory caspases may play a role together with caspase-1 to promote IFN-γ-mediated inflammasome responses. Collectively, our data indicate that caspase-1, caspase-4, and caspase-5 participate in inflammasome responses to *L. pneumophila* infection in IFN-γ-primed hMDMs.

**Fig. 2.**
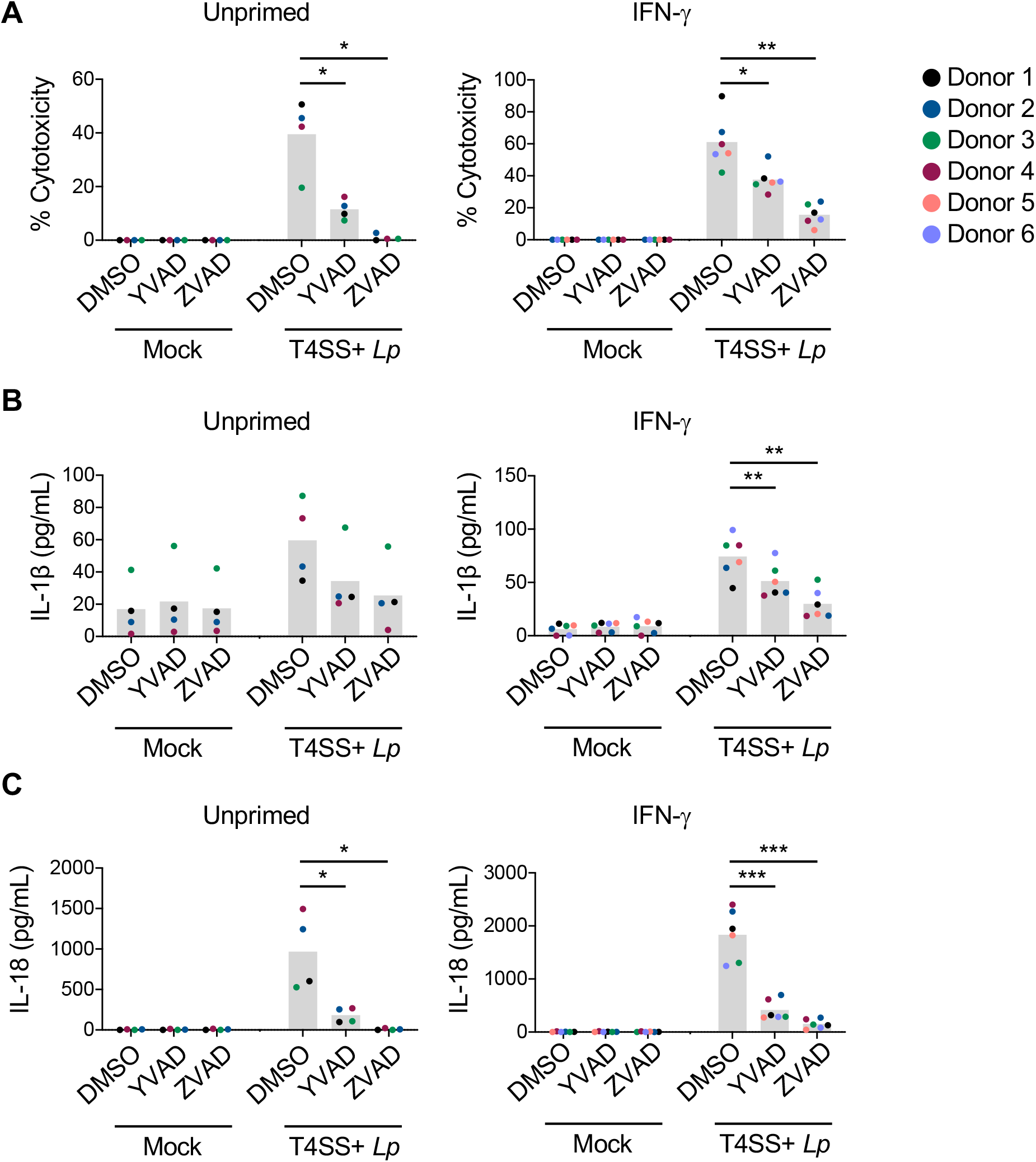
Caspase-1 and additional caspases promote inflammasome activation in response to *L. pneumophila*. (A, B, C) Primary hMDMs were left unprimed or primed with IFN-γ (100 U/mL) overnight and treated with the inhibitors YVAD or ZVAD, or DMSO control for one hour followed by infection with T4SS+ *Lp* for four hours. (A) Cell death was measured using lactate dehydrogenase release assay and normalized to mock-infected cells. (B and C) IL-1β and IL-18 levels in the supernatant were measured by ELISA. Shown are the pooled results of four to six independent experiments using hMDMs from different healthy human donors. Each data point represents the mean of triplicate infected wells from an individual donor. *P< 0.05, **P< 0.01, and ***P< 0.001 by paired t-test.

### IFN-γ upregulates human GBPs

IFN-γ induces expression of a large number of genes that contribute to antimicrobial defense. In mice, two IFN-inducible gene families that promote inflammasome activation in macrophages are the GBPs and IRGs. Their assigned functions include binding and rupturing the phagosome of vacuolar pathogens, as well as directly lysing bacteria that escape the phagosome and enter the cytosol (20-22, 52). These activities lead to release of pathogen-derived products such as lipopolysaccharide (LPS) and DNA into the cytosol, resulting in downstream inflammasome activation. Mice have 11 GBPs and 23 IRGs, whereas humans have seven GBPs and two IRG genes (53). Human GBPs, like their murine counterparts, are IFN-inducible, whereas human IRGs are not induced by IFN stimulation (37, 53, 54).

Thus, we chose to test whether human GBPs might play a role in the enhanced inflammasome responses of IFN-γ-primed cells to *L. pneumophila*. We first asked whether GBP expression is upregulated by IFN-γ in THP-1-derived macrophages and hMDMs. In THP-1 cells, we found that expression of all GBPs was induced in response to IFN-γ, and *GBP1-5* mRNA levels were significantly upregulated in hMDMs following IFN-γ treatment (Fig. 3A and B). Following IFN-γ-priming, we observed high relative expression of *GBP1, GBP2, GBP3, GBP4*, and *GBP5*, whereas there was very low relative expression of *GBP6* and *GBP7* in THP-1 cells (Fig. S2A) and hMDMs (Fig. S2B), in agreement with previous findings (37). Furthermore, priming hMDMs with increasing amounts of IFN-γ led to a dose-dependent increase in GBP mRNA levels (Fig. 3C and S2C). Protein levels of GBP1, GBP2, GBP4, and GBP5 were also increased in a dose-dependent manner in response to IFN-γ (Fig. 3D). Thus, human GBPs are transcriptionally and translationally induced by IFN-γ in macrophages, in agreement with previous findings (37, 54).

**Fig. 3.**
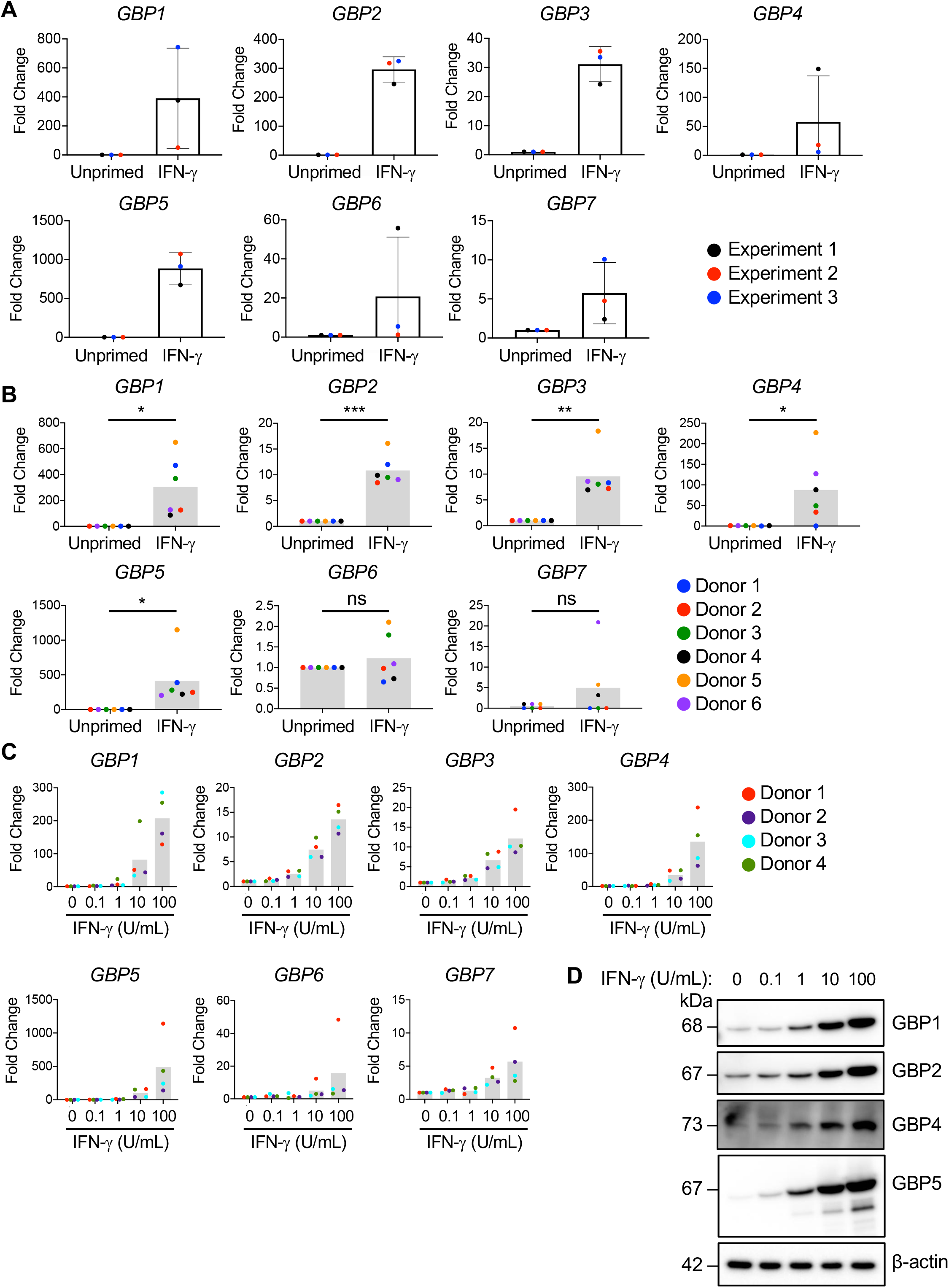
Human GBPs are transcriptionally and translationally upregulated in response to IFN-γ. PMA-differentiated THP-1 cells (A) or primary hMDMs (B) were either left unprimed or primed with IFN-γ (100 u/mL) for 18 or 20 hours, respectively. (C and D) hMDMs were left unprimed or primed with IFN-γ at the indicated concentrations for 20 hours. (A, B, C) Transcript levels of *GBP1-7* were determined by quantitative RT-PCR and fold change was calculated by normalizing to the housekeeping gene HPRT for each sample and then to the unprimed sample. Shown are the pooled results of three independent experiments (A) or six independent experiments using hMDMs different healthy human donors (B), with each data point representing the value for each experiment (A) or an individual donor (B). *P< 0.05, **P< 0.01, and ***P< 0.001 by paired t-test. (C) Shown are the pooled results of four independent experiments using hMDMs from different healthy human donors and each data point represents the value of an individual donor. (D) Immunoblot analysis was conducted on lysates for GBP1, GBP2, GBP4, GBP5, and β-actin. Western blot is representative of four independent experiments using hMDMs from different healthy human donors.

### Human GBP1 contributes to maximal IFN-γ-dependent inflammasome responses to *L. pneumophila*

Since GBP1-5 were significantly upregulated in hMDMs, we next wanted to test whether these GBPs play a role in human inflammasome responses to *L. pneumophila*. We therefore individually silenced expression of *GBP1-5* prior to IFN-γ treatment and T4SS+ *Lp* infection in hMDMs. Notably, specific knockdown of *GBP1* significantly decreased cell death and IL-1β and IL-18 secretion following *L. pneumophila* infection in IFN-γ-primed hMDMs, indicating that GBP1 plays a non-redundant role in inflammasome responses against *L. pneumophila* infection (Fig. 4A and B). GBP3 knockdown resulted in significantly decreased IL-1β release but did not affect cell death or IL-18 release. In contrast, knockdown with siRNAs against *GBP2, 4*, and 5 did not decrease cell death or cytokine secretion. Importantly, we examined the knockdown efficiencies for hMDMs treated with siRNA for each GBP and found that siRNA knockdown was specific for each GBP and did not affect the expression levels of the remaining GBPs (Fig. 4C). Collectively, these data indicate that human GBP1 is important for promoting maximal cell death and IL-1 family cytokine release.

**Fig. 4.**
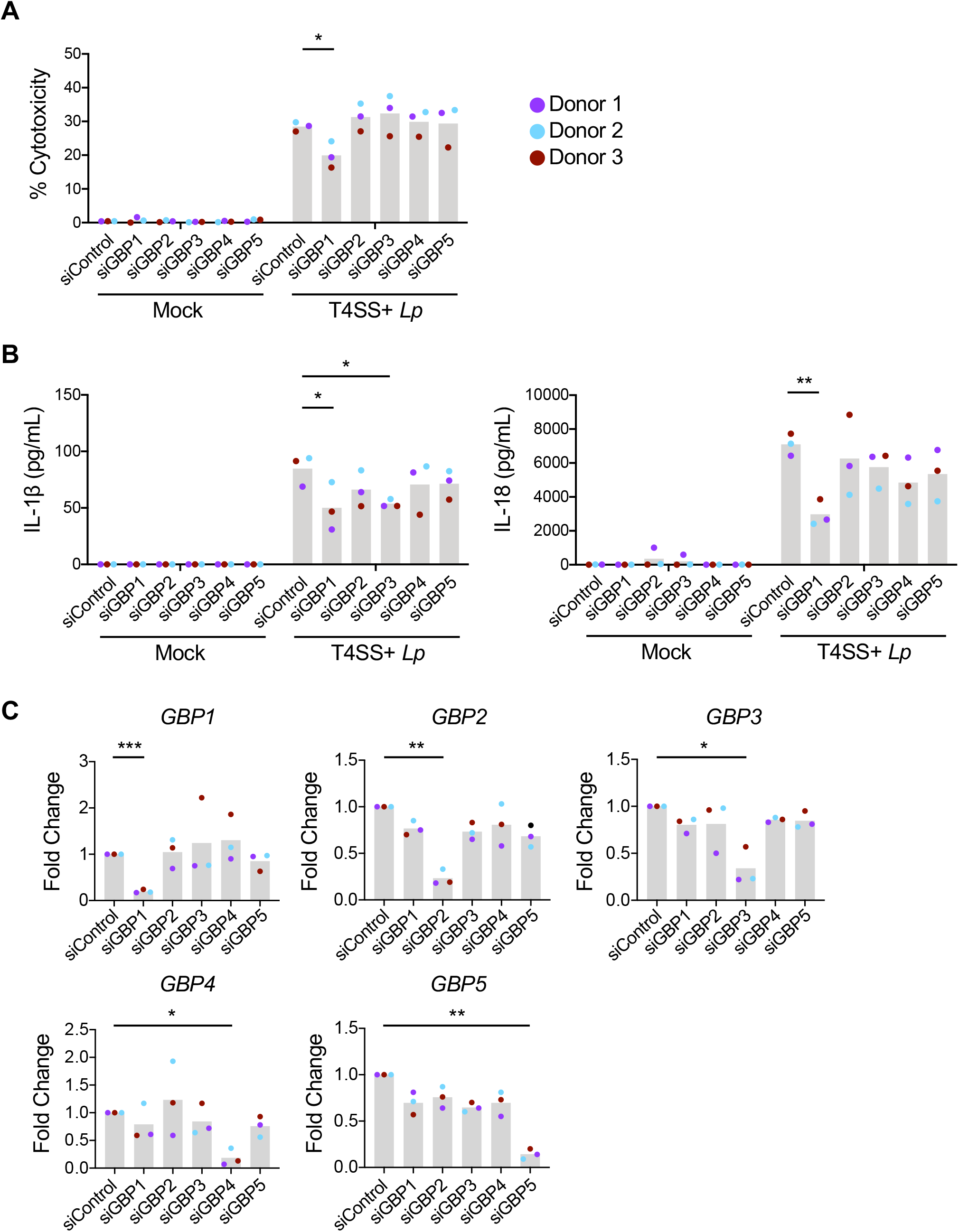
GBP1 is required for maximal inflammasome activation. Primary hMDMs were transfected with 30nM siRNA specific for individual GBP or scrambled control siRNA (siControl), primed with IFN-γ (100 U/mL) overnight, and infected with T4SS+ *Lp* for four hours. (A) Cell death was measured using lactate dehydrogenase release assay and normalized to mock infected cells. (B) IL-1β and IL-18 levels in the supernatant were measured by ELISA. (C) Transcript levels of *GBP1-5* in ‘mock’ samples were determined by quantitative RT-PCR and fold change was calculated by normalizing to the housekeeping gene HPRT for each sample and then to the siControl sample. (A, B, C) Shown are the pooled results of three independent experiments using hMDMs from different healthy human donors. (A and B) Each data point represents the mean of triplicate infected wells from an individual donor. *P< 0.05, **P< 0.01, and ***P< 0.001 by paired t-test. (C) Each data point represents the value of an individual donor.

### IFN-γ promotes GBP localization to *L. pneumophila* in a T4SS-dependent manner

Since our data indicated that human GBP1 is required for maximal inflammasome activation during infection with *L. pneumophila*, we next wanted to elucidate how GBP1 could be promoting this response. Mouse Gbp2 colocalizes to the *Salmonella*-containing vacuole (SCV) while its predicted human ortholog, GBP1, colocalizes with *S*. Typhimurium in macrophages (21, 36). Nevertheless, whether human GBP1 disrupts the SCV or the *Salmonella* outer membrane remains unknown. We hypothesized that human GBP1 might play a similar role in *L. pneumophila* infection and would be predicted to colocalize with the LCV in IFN-γ-primed macrophages. To test this hypothesis, we infected IFN-γ-primed and unprimed hMDMs with dsRED-expressing T4SS+ *Lp* and stained for GBP1. While there was little to no GBP1 expression or colocalization with *L. pneumophila* in unprimed cells, there was a significant increase in the percentage of infected cells containing GBP1-positive *L. pneumophila* following IFN-γ priming (Fig. 5A and B). Approximately 60% of infected cells contained *L. pneumophila* that colocalized with GBP1. In contrast, GBP1 was distributed throughout the cytoplasm in uninfected IFN-γ-primed hMDMs (Fig. S3A). The secondary antibodies used for anti-GBP1 staining did not associate with *L. pneumophila* when used alone and only stained cells when primary anti-GBP1 antibodies were used (Fig. S3B). These data indicate that GBP1 is recruited to *L. pneumophila* and/or the LCV within IFN-γ-primed hMDMs.

**Fig. 5.**
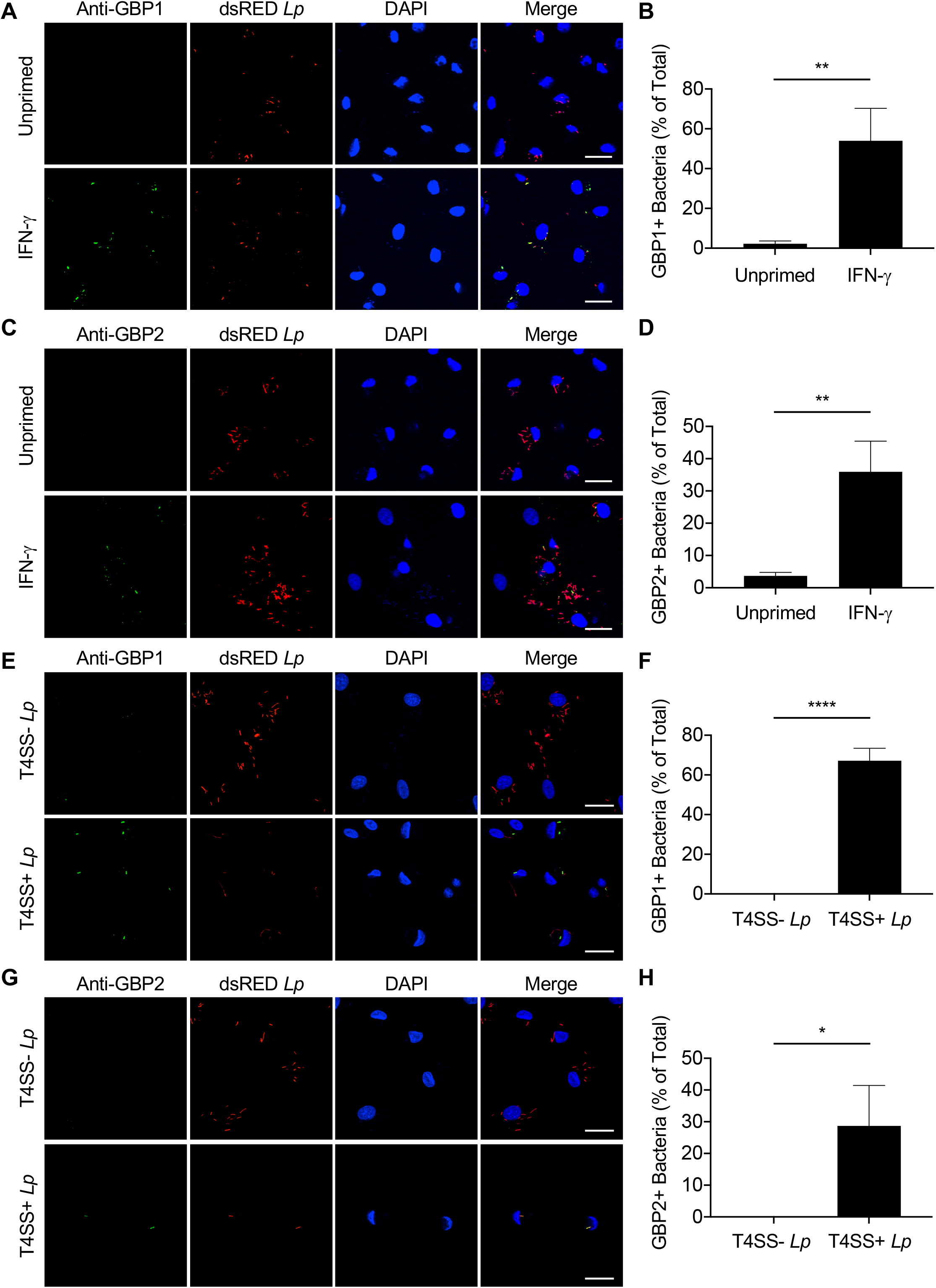
IFN-γ promotes the colocalization of GBP1 and GBP2 with *L. pneumophila* in a T4SS-dependent manner. (A-D) Primary hMDMs were either left unprimed or primed with IFN-γ (100 U/mL) overnight and infected with dsRED-expressing T4SS+ *Lp* for two hours. Representative fluorescence micrographs of anti-GBP1 (A) or anti-GBP2 (C) antibody staining in dsRED-T4SS+ *Lp*-infected hMDMs and quantification of the percentage of hMDMs containing GBP1+ *Lp* (B) or GBP2+ *Lp* (D) out of total infected hMDMs. Graphs show the mean and s.d. of technical triplicates and data are representative of three independent experiments using hMDMs from different healthy human donors. **P< 0.01 by unpaired t-test. (E-H) Primary hMDMs were primed with IFN-γ (100 U/mL) overnight and infected with dsRED-expressing T4SS-*Lp* or T4SS+ *Lp* for two hours. Representative fluorescence micrographs of anti-GBP1 (E) or anti-GBP2 (G) antibody staining in dsRED-T4SS- or dsRED-T4SS+ *Lp*-infected hMDMs and quantification of the percentage of hMDMs containing GBP1+ *Lp* (F) or GBP2+ *Lp* (H) out of total infected hMDMs. Graphs show the mean and s.d. of technical triplicates and data are representative of two independent experiments using hMDMs from different healthy human donors. *P< 0.05 and ****P<0.0001 by unpaired t-test.

While it is unclear whether GBP1 binds to the LCV or the bacterial outer membrane, human GBP1 does bind to the outer membrane of the cytosolic bacterium, *S. flexneri*, and additional GBPs are also recruited to inhibit its actin motility (34, 35). Thus, we tested whether GBP2 also localized to *L. pneumophila*. We also observed a significantly increased percentage of hMDMs harboring GBP2+ *L. pneumophila* following IFN-γ priming compared to unprimed cells (Fig. 5C and D), although to a lower extent compared to GBP1+ *L. pneumophila*. Furthermore, secondary antibodies used for anti-GBP2 staining did not stain when used alone and only colocalized with *L. pneumophila* when primary anti-GBP2 antibodies were used (Fig. S3C). Collectively, these findings show that both GBP1 and GBP2 are recruited to *L. pneumophila* and/or the LCV in IFN-γ-primed hMDMs.

In mouse macrophages, colocalization of GBPs with the LCV is dependent on the T4SS, while GBP colocalization with the *Yersinia*-containing vacuole requires the presence of type III secretion system translocon components (26, 55). These findings indicate that murine GBPs respond to secretion systems that are key signatures of bacterial virulence. However, whether human GBPs also detect PCVs that contain bacteria expressing virulence-associated secretion systems is unclear. Notably, only T4SS+ *Lp*, but not T4SS-*Lp*, exhibited robust colocalization with GBP1 and GBP2 in IFN-γ-primed hMDMs (Fig. 5E-H). Collectively, these data suggest that GBP1 and GBP2 are upregulated in response to IFN-γ priming and following infection, are recruited to *L. pneumophila* in a T4SS-dependent manner.

### IFN-γ and GBP1 promote the rupture of LCVs

We next wanted to determine how IFN-γ and GBP1 promote increased inflammasome activation during *L. pneumophila* infection. We first tested whether IFN-γ treatment results in an increase of ruptured LCVs, which would allow *L. pneumophila* to become more accessible for recognition by cytosolic inflammasome sensors. We utilized a differential permeabilization assay to distinguish between vacuolar and cytosolic *L. pneumophila* in the presence and absence of IFN-γ priming (56). We compared unprimed and IFN-γ-primed hMDMs that were infected with dsRED-expressing T4SS+ *Lp* and then treated with the detergent digitonin, which selectively permeabilizes the plasma membrane while leaving intracellular membranes intact. The cells were then immunostained with an antibody for *L. pneumophila*, followed by staining with an Alexa 488-labeled secondary antibody that fluoresces green. Thus, dsRED-expressing *L. pneumophila* contained within an intact vacuole only fluoresce red, while dsRED-expressing *L. pneumophila* within a ruptured vacuole will fluoresce both green and red (Fig. 6A). We found that a significantly increased percentage of hMDMs primed with IFN-γ contained *L. pneumophila* that stained with anti-*L. pneumophila* antibody and fluoresced green compared to unprimed cells (Fig. 6B and C). Treatment with the detergent saponin, which permeabilizes all cell membranes, resulted in similar percentages of unprimed and IFN-γ-primed hMDMs containing bacteria that were stained by anti-*L. pneumophila* antibody (Fig S4A and B). The secondary antibody stained only in the presence of anti-*L. pneumophila* antibody (Fig S4C), indicating that the signal we observed in digitonin-permeabilized, IFN-γ-primed hMDMs was indeed due to an increased presence of cytosolic *L. pneumophila*. These results indicate that IFN-inducible host factors promote rupture of the LCV, resulting in increased *L. pneumophila* exposure to the host cell cytosol.

**Fig. 6.**
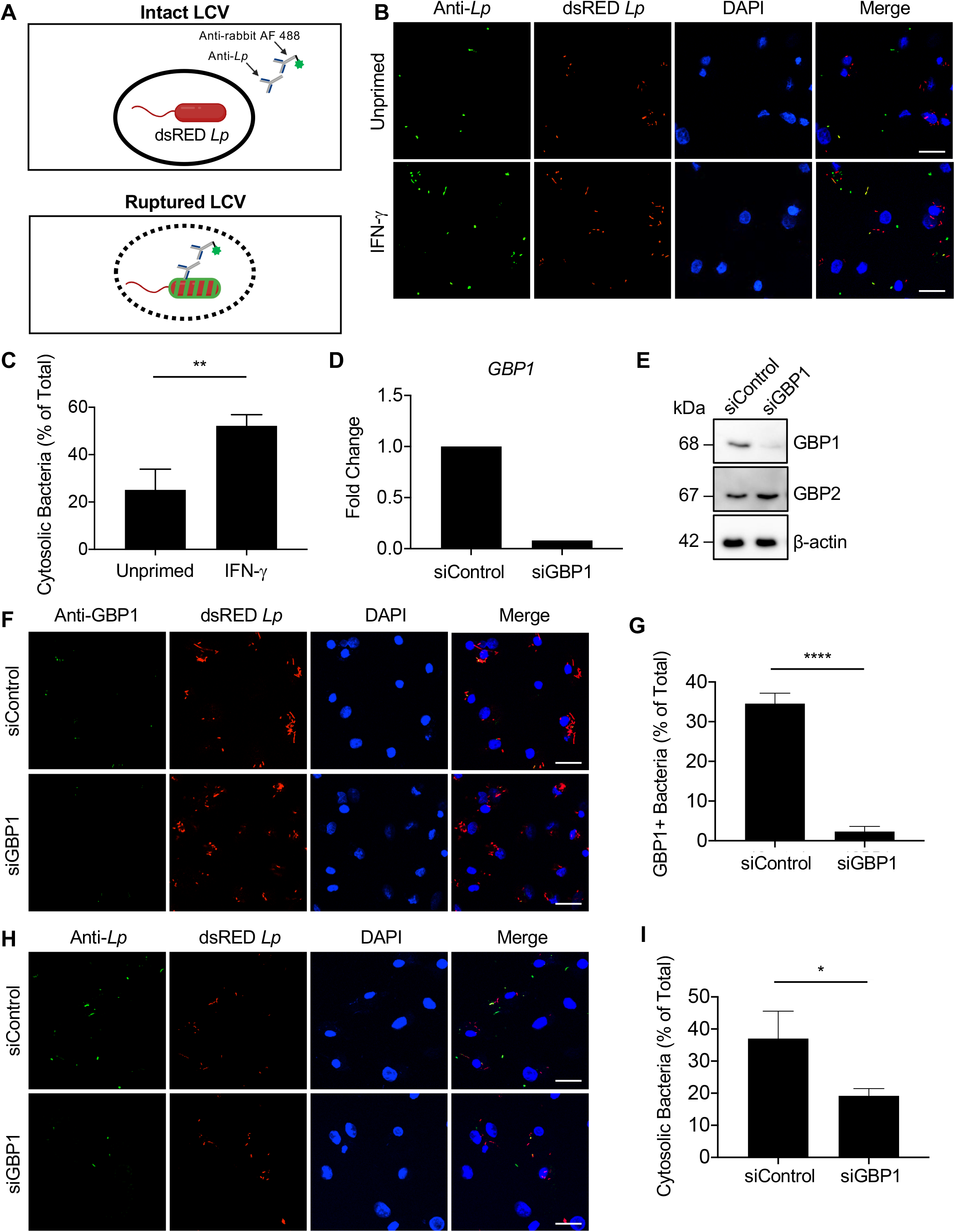
IFN-γ and GBP1 promote the rupture of LCVs in hMDMs. (A) Schematic of vacuolar *Lp*, which fluoresces red, and cytosolic *Lp*, which is stained green and fluoresces red. (B and C) Primary hMDMs were either left unprimed or primed with IFN-γ (100 U/mL) overnight and infected with dsRED-expressing T4SS+ *Lp* for two hours. (B) Representative fluorescence micrographs of anti-*Lp* antibody staining followed by Alexa 488-conjugated secondary antibody staining in digitonin-permeabilized dsRED-T4SS+ *Lp*-infected hMDMs. (C) Quantification of the percentage of hMDMs harboring cytosolic *Lp* out of total infected hMDMs. (D-I) Primary hMDMs were transfected with 5 pmol siRNA specific for GBP1 (siGBP1) or scrambled control siRNA (siControl) for at least 48 h, primed with IFN-γ (100 U/mL) overnight, and infected with dsRED-expressing T4SS+ *Lp* for two hours. (D) *GBP1* transcript levels in ‘mock’ samples were determined by quantitative RT-PCR. Fold change was calculated by normalizing to the housekeeping gene *HPRT* and then to the siControl sample. (E) Immunoblot analysis was conducted on ‘mock’ lysates for GBP1, GBP2, and β-actin. (F) Representative fluorescence micrographs of anti-GBP1 antibody staining in dsRED-T4SS+ *Lp*-infected hMDMs. (G) Quantification of the percentage of hMDMs containing GBP1+ *Lp* out of total infected hMDMs. (H) Representative fluorescence micrographs of anti-*Lp* antibody staining followed by Alexa 488-conjugated secondary antibody staining in digitonin-permeabilized dsRED-T4SS+ *Lp*-infected hMDMs. (I) Quantification of the percentage of hMDMs harboring cytosolic *Lp* out of total infected hMDMs. Graphs show the mean and s.d. of technical triplicates and data are representative of three independent experiments using hMDMs from different healthy human donors. *P<0.05, **P< 0.01 and ****P<0.0001 by unpaired t-test. (D and E) Data are representative of three independent experiments using hMDMs from different healthy human donors.

Since GBP1 colocalizes with *L. pneumophila* (Fig. 5) and GBP1 is important for maximal inflammasome responses to *L. pneumophila* in IFN-γ-primed hMDMs (Fig. 4), we hypothesized that GBP1 might contribute to the disruption of LCV integrity. Therefore, we conducted the phagosome integrity assay in *GBP1*-silenced IFN-γ-primed hMDMs. We confirmed efficient and specific *GBP1* knockdown at the mRNA and protein levels compared to control siRNA treatment (Fig. 6D and E). Additionally, a significantly lower percentage of infected hMDMs treated with *GBP1* siRNA contained GBP1+ *L. pneumophila* compared to control siRNA-treated hMDMs (Fig. 6F and G). Interestingly, a significantly decreased percentage of *GBP1* siRNA-treated hMDMs contained bacteria that were stained by anti-*L. pneumophila* antibody following digitonin permeabilization compared to hMDMs treated with control siRNA (Fig. 6H and I), indicating that there is a significant decrease in the percentage of cells containing ruptured LCVs following *GBP1* knockdown. In contrast, following saponin permeabilization of all cellular membranes, a similar percentage of hMDMs contained bacteria that stained positive for anti-*L. pneumophila* antibody following control or GBP1 siRNA treatment (Fig. S4E and F), whereas staining with secondary antibody alone revealed negligible background staining (Fig. S4D and G). Collectively, these findings indicate that GBP1 plays a key role in IFN-γ-dependent disruption of the LCV in primary human macrophages, thus allowing for increased access of *L. pneumophila* to the host cell cytosol.

## Discussion

Our data reveal that human GBP1 is crucial for robust inflammasome activation in response to *L. pneumophila* infection in IFN-γ-primed primary human macrophages. These findings are the first to report the role of human GBPs in inflammasome activation in response to *L. pneumophila* infection. We show that IFN-γ leads to enhanced cell death and proinflammatory cytokine release during *L. pneumophila* infection and that this inflammasome response involves caspase-1, capsase-4, caspase-5, and GSDMD processing. We also find that GBP1 colocalizes with *L. pneumophila* in a T4SS-dependent manner and promotes increased access of *L. pneumophila* to the host cell cytosol, indicating that GBP1 facilitates disruption of the LCV. Our findings suggest a model in which human GBP1 promotes the liberation of *L. pneumophila* components into the host cell cytosol to allow for increased inflammasome sensing and activation. Intriguingly, murine GBPs do not disrupt the LCV, but rather promote outer membrane disruption of cytosolic *L. pneumophila* (22). Together, these findings suggest that human and murine GBPs play distinct roles in mediating inflammasome responses against *L. pneumophila*.

Although mice encode 11 GBPs and humans encode seven GBPs, there are some GBPs shared between mice and humans, with mouse Gbp2 and Gbp5 thought to be the orthologs of human GBP1 and GBP5, respectively (33). These murine orthologs may provide insight into the functions of human GBPs, since most experimental studies aimed at elucidating GBP functions have been conducted in mice. Mouse GBPs colocalize with vacuoles that harbor bacterial secretion systems or bacterial translocon components (26, 55). Mouse Gbp2 promotes lysis of the SCV and activation of the noncanonical inflammasome, while its ortholog human GBP1 colocalizes with *S*. Typhimurium and promotes caspase-4-mediated pyroptosis (21, 36). In contrast to *S*. Typhimurium, mouse GBPs do not mediate vacuole disruption for other bacterial pathogens, but instead facilitate lysis of cytosolic bacteria (20–22). Whether human GBP1 is recruited to pathogen-containing vacuoles and whether it promotes lysis of pathogen-containing vacuoles or bacteria was unknown. Importantly, our findings reveal that GBP1 targets the LCV in a T4SS-dependent manner and furthermore, that GBP1 promotes vacuolar disruption and increased exposure of *L. pneumophila* to the host cell cytosol. Thus, human and mouse orthologs may have both distinct and overlapping functions. Additional studies will further elucidate the roles of human GBPs in response to other bacterial infections.

Our data show that human GBP1 and GBP2 colocalize with *L. pneumophila* in a T4SS-dependent manner, but whether and how these GBPs are recruited and bound to the LCV and/or bacterial outer membrane still remains to be determined. Mouse Gbp2 colocalizes with pathogen-containing vacuoles containing bacterial secretion systems in a galectin-3-dependent manner (55). Whether galectins facilitate human GBP1 recruitment to pathogen-containing vacuoles is unknown. Furthermore, human and mouse GBP1, GBP2, and GBP5 have a C-terminal CaaX prenylation motif that facilitates membrane binding and oligomerization with other GBPs (57). Human GBP1 binds to the outer membrane of *S. flexneri* and colocalizes with *S*. Typhimurium in a manner dependent on its isoprenylation and GTPase activity (34–36). In addition, human GBP1 colocalizes with a *S. flexneri* mutant lacking the O-antigen less frequently than with the wild-type strain, indicating that host recognition of O-antigen enables GBP1 binding to *S. flexneri* (35). It would be of interest to determine whether the CaaX motif in human GBP1 and GBP2 are necessary for colocalization with *L. pneumophila* and what bacterial or vacuolar components they are binding to. Although we found that GBP2 colocalized with *L. pneumophila*, siRNA-mediated silencing of GBP2 did not have an effect on inflammasome activation. It is possible that GBP2 is not required for inflammasome responses to *L. pneumophila* or that siRNA-mediated knockdown in primary hMDMs was not efficient enough to reveal a role for GBP2. Further studies will discern between these possibilities. Since we found that GBP1 promotes inflammasome activation, it would also be of interest to determine whether GBP1 may act as an initiator GBP that recruits additional GBPs, similar to what has been observed in with *S. flexneri* (34, 35, 58), and whether there is a synergistic role for human GBPs.

Inflammasome activation is triggered in response to sensing of bacterial products within the cytosol. Vacuolar localization of *L. pneumophila* within its ER-derived vacuole would presumably limit the ability of host cells to recognize *L. pneumophila* components. However, when the integrity of the LCV is compromised, either by host factors or in the case of bacterial mutants that cannot maintain vacuolar integrity *L. pneumophila* becomes more accessible for recognition by host cytosolic sensors (22). We show that IFN-γ priming in primary human macrophages results in an increased frequency of ruptured LCVs, indicating that IFN-inducible host cell factors promote disruption of the LCV. Our data indicate that GBP1 is one such factor. While we cannot formally conclude that GBP1-mediated rupture of the LCV is the proximal cause of downstream inflammasome activation, this rupture likely results in increased exposure of *L. pneumophila* products to the host cell cytosol, thus making the bacteria vulnerable to inflammasome sensing. Human GBPs may also target and promote destabilization of the outer membrane of *L. pneumophila* to enable the release of bacterial components, including LPS and DNA, for inflammasome sensing. Murine GBPs encoded on chromosome 3 promote the disruption of the outer membrane of the cytosolic *L. pneumophila* mutant lacking the effector SdhA, which is important for maintaining the vacuole integrity of the LCV (22). Mouse macrophages lacking chromosome 3 GBPs that were infected with the *ΔsdhA* mutant showed a decrease in pyroptosis and release of DNA into the cytosol, indicating that one or more chromosome 3 GBPs contribute to inflammasome activation in response to cytosolic bacteria. Since mouse GBPs mediate the disruption of cytosolic *L. pneumophila*, it is possible that human GBP1 or other GBPs may also enable disruption of the *L. pneumophila* outer membrane to release bacterial components that subsequently lead to inflammasome activation.

Overall, our findings reveal a critical role for IFN-γ and human GBP1 in promoting human inflammasome responses against *L. pneumophila*. In particular, our study illuminates a key function for human GBP1 in disrupting the pathogen-containing vacuole. These findings indicate that human GBPs have distinct roles compared to mouse GBPs in promoting inflammasome responses to *L. pneumophila* and provide insight into human cell-autonomous responses to a vacuolar bacterial pathogen.

## Materials and Methods

### Ethics statement

All studies on primary human monocyte-derived macrophages (hMDMs) were performed in compliance with the requirements of the US Department of Health and Human Services and the principles expressed in the Declaration of Helsinki. Samples obtained from the University of Pennsylvania Human Immunology Core are considered to be a secondary use of deidentified human specimens and are exempt via Title 55 Part 46, Subpart A of 46.101 (b) of the Code of Federal Regulations.

### Cell culture

THP-1 cells (TIB-202; American Type Culture Collection) were maintained in RPMI supplemented with 10% (vol/vol) heat-inactivated FBS, 0.05 nM β-mercaptoethanol, 100 IU/mL penicillin, and 100 μg/mL streptomycin at 37°C in a humidified incubator. The day before stimulation, cells were replated in media without antibiotics in a 48-well plate at a concentration of 2 × 10^5^ cells per well or in a 96-well plate at a concentration of 1 × 10^5^ cells per well and incubated with phorbol 12-myristate 13-acetate (PMA) for 24 hours to allow differentiation into macrophages. Media was replaced with RPMI without serum for infections in 48-well plate.

Primary human monocytes from deidentified healthy human donors were obtained from the University of Pennsylvania Human Immunology Core. Monocytes were cultured in RPMI supplemented with 10% (vol/vol) heat-inactivated FBS, 2 mM L-glutamine, 100 IU/mL penicillin, 100 μg/mL streptomycin, and 50 ng/mL recombinant human M-CSF (Gemini Bio Products). Cells were cultured for 4 days in 10 mL of media in 10 cm-dishes at 4-5 × 10^5^ cells/mL, followed by addition of 10 mL of fresh growth media for an additional 2 days for complete differentiation into macrophages. The day before macrophage stimulation, cells were rinsed with cold PBS, gently detached with trypsin-EDTA (0.05%) and replated in media without antibiotics and with 25 ng/mL M-CSF in a 48-well plate at a concentration of 1 × 10^5^ cells per well or in a 24-well plate at a concentration of 2 × 10^5^ cells per well.

### Macrophage stimulation

In infection experiments, PMA-differentiated THP-1 cells and primary human monocyte-derived macrophages (hMDMs) were either left unprimed or were primed overnight with recombinant human IFN-γ (R&D Systems) at a concentration of 100 U/mL. In dose-response experiments, hMDMs were either left unprimed or primed with 0.1, 1, 10, or 100 U/mL of IFN-γ for 20 hours.

### Bacterial strains and macrophage infection

All *Legionella pneumophila* infections used strains derived from the serogroup 1 clinical isolate Philadelphia-1. Where indicated, strains utilized were derived from the Lp02 strain (*rpsL, hsdR, thyA*), which is a thymidine auxotroph. The isogenic Lp02 (*rpsL, hsdR, thyA*) flagellin mutant, Δ*flaA* (T4SS+ *Lp*), and avirulent *dotA* mutant, Lp03 (T4SS-*Lp*), which are both thymidine auxotrophs, were used to infect PMA-differentiated THP-1 cells and primary hMDMs (39, 41, 59). Δ*flaA* (T4SS+) or Δ*dotA* (T4SS-) *L. pneumophila* strains on the JR32 background (*rpsL, hsdR*) carrying pSW001, which allows for constitutive dsRED expression, were used in immunofluorescence experiments (60, 61). All *L. pneumophila* strains were grown as a stationary patch for 48 hours on charcoal yeast extract agar plates at 37°C (62). Bacteria were resuspended in PBS and added to the cells at a multiplicity of infection (MOI) of 10 in 48-well and 24-well plate experiments. Infected cells were then centrifuged at 290 × g for 10 min and incubated at 37°C. For immunofluorescence experiments, cells were infected for 2 hours. For all additional infection experiments, cells were infected for 4 hours. For all experiments, mock-infected cells were treated with PBS.

### Caspase inhibitor treatments

25 μM of caspase-1 inhibitor Ac-YVAD-cmk (Sigma-Aldrich SML0429) and 20 μM of pan-caspase inhibitor Z-VAD(OMe)-FMK (SM Biochemicals SMFMK001) were added to primary hMDMs 1 hour before infection.

### siRNA-mediated knockdown

All of the Silencer Select siRNA oligos targeting human *GBP* mRNA were purchased from Thermo Fisher Scientific. Individual siRNA targeting *GBP1* (s5620), *GBP2* (s5623), *GBP3* (5628), *GBP4* (s41805), and *GBP5* (s41810) were used. The two Silencer Select negative control siRNAs (Silencer Select Negative Control No. 1 siRNA and Silencer Select Negative Control No. 2 siRNA) were purchased from Life Technologies (Ambion). In experiments where *GBP1-5* were individually knocked down, primary hMDMs were replated in media without antibiotics in a 48-well plate, as described above, three days before infection. Two days before infection, 30 nM of total siRNA were transfected into macrophages using HiPerFect transfection reagent (Qiagen) following the manufacturer’s protocol. 16 hours before infection, media was replaced with fresh antibiotic-free media containing 100 U/mL IFN-γ. In immunofluorescence experiments where *GBP1* was knocked down, primary hMDMs were replated in media without antibiotics on glass coverslips in a 24-well plate as described above four days before infection. Three days before infection, 5 pmol of total siRNA were transfected into macrophages using Lipofectamine RNAiMAX transfection reagent (Thermo Fisher Scientific) following the manufacturer’s protocol. 16 hours before infection, media was replaced with fresh antibiotic-free media containing 100 U/mL IFN-γ.

### Quantitative RT-PCR Analysis

RNA was isolated using the RNeasy Plus Mini Kit (Qiagen) following the manufacturer’s protocol. Cells were lysed in 350 μL RLT buffer with β-mercaptoethanol and centrifuged through a QIAshredder spin column (Qiagen). cDNA was synthesized from isolated RNA using SuperScript II Reverse Transcriptase (Invitrogen) following the manufacturer’s protocol. Quantitative PCR was conducted with the CFX96 real-time system from Bio-Rad using the SsoFast EvaGreen Supermix with Low ROX (Bio-Rad). Transcript levels for each gene were normalized to the housekeeping gene HPRT for each sample, and samples were normalized to unprimed sample or to control siRNA-treated sample using the 2^−ΔΔCt^ (cycle threshold) method to calculate fold change. Relative expression was calculated by normalizing gene-specific transcript levels to HPRT transcript levels for each sample using the 2^−ΔCt^ method. Primer sequences from primer bank used for *HPRT1*, *GBP1-6*, *CASP4*, and *CASP5* or from Lagrange, et al. for *GBP7* are in Supplementary Table 1.

### LDH cytotoxicity assay

Macrophages were infected in a 48-well plate as described above and harvested supernatants were assayed for cell death by measuring loss of cellular membrane integrity via lactate dehydrogenase (LDH) activity. LDH release was quantified using an LDH Cytotoxicity Detection Kit (Clontech) according to the manufacturer’s instructions and normalized to mock-infected cells.

### Real-time propidium iodide uptake assay

To measure live kinetics of cell membrane permeability, THP-1 cells were plated as described above in a black, flat-bottom 96-well plate (Cellstar), primed with 100 U/mL IFN-γ for 24 hours, and infected with T4SS+ *Lp* at an MOI of 50 in media containing 1X HBSS without phenol red, 20 mM HEPES, and 10% (vol/vol) heat-inactivated FBS. Infected cells were centrifuged at 290 × g for 10 min. The cells were supplemented with 5 μM propidium iodide (PI, P3566, Invitrogen) and incubated for 10 min at 37°C to allow the cells to equilibrate. Then, the plate was sealed with adhesive optical plate sealing film (Microseal, Bio-Rad) and placed in a Synergy H1 microplate reader (BioTek) pre-heated to 37°C. PI fluorescence was measured every hour for 4 hours.

### ELISA

Macrophages were infected in a 48-well plate as described above and harvested supernatants were assayed for cytokine levels using ELISA kits for human IL-1β (BD Biosciences) and IL-18 (R&D Systems).

### Immunoblot analysis

In experiments where macrophages were plated in a 48-well plate, cells were lysed in 1× SDS/PAGE sample buffer, and low-volume supernatants (90 μL media per well of a 48-well plate) were mixed 1:1 with 2× SDS/PAGE sample buffer containing Complete Mini EDTA-free Protease Inhibitor Mixture (Roche). In experiments where primary hMDMs were plated in a 24-well plate and infected with T4SS-*Lp*, T4SS+ *Lp*, or mock infected with PBS, cells were lysed in 1X SDS/PAGE sample buffer, and supernatants were treated with trichloroacetic acid (TCA) overnight at 4°C and centrifuged at maximum speed for 15 min. Precipitated supernatant pellets were washed with ice-cold acetone, centrifuged at maximum speed for 10 min, and resuspended in 1X SDS/PAGE sample buffer. Protein samples were boiled for 5 min, separated by SDS/PAGE on a 12% (vol/vol) acrylamide gel, and transferred to PVDF Immobilon-P membranes (Millipore). Primary antibodies specific for human IL-1β (clone 8516; R&D Systems), caspase-1 (2225S; Cell Signaling), caspase-4 (4450S; Cell Signaling), caspase-5 (D3G4W; 46680S; Cell Signaling), Gasdermin-D (126-138; G7422; Sigma-Aldrich), GBP1 (ab131255, Abcam), GBP2 (sc-271568, Santa Cruz), GBP4 (17746-1-AP, Proteintech), GBP5 (D3A5O, 67798S; Cell Signaling) and β-actin (4967L; Cell Signaling) were used. HRP-conjugated secondary antibodies anti-rabbit IgG (7074S; Cell Signaling) and anti-mouse IgG (7076S; Cell Signaling) were used. For detection, ECL Western Blotting Substrate or SuperSignal West Femto (both from Pierce Thermo Scientific) were used as the HRP substrate.

### Immunofluorescence microscopy

Primary hMDMs were plated on glass coverslips in a 24-well plate as described above. After 2 hours of infection with dsRED-*Lp*, cells were washed 2 times with PBS and fixed with 4% paraformaldehyde for 10 min at 37°C. Following fixation, cells were washed and permeabilized with 0.2% Triton X-100 for 10 min. Cells were washed, blocked with 10% BSA for 1 hour, and stained with primary antibodies (identified below) for 1 hour. Cells were washed with PBS and incubated with the appropriate Alexa-Fluor-conjugated secondary antibodies (identified below) for 1 hour, followed by washes and mounted on glass slides with DAPI mounting medium (Sigma Fluoroshield). Primary antibodies used were rabbit anti-GBP1 (1:100 dilution; Abcam) and mouse anti-GBP2 (1:50 dilution; Santa Cruz). Secondary antibodies used at a dilution of 1:4000 were goat anti-rabbit conjugated to Alexa Fluor 488 (4412S; Cell Signaling) and goat anti-mouse conjugated to Alexa Fluor 488 (A11029; Life Technologies). Coverslips were imaged on a Leica SP5 FLIM confocal microscope at a magnification of 63× and the percentage of infected cells containing GBP1+ or GBP2+ intracellular bacteria out of the total number of infected cells were quantified.

### Phagosome integrity assay

The phagosome integrity assay was performed as previously published (56), with some modifications. To distinguish between cytosolic and vacuolar bacteria, primary hMDMs were plated on glass coverslips in a 24-well plate as described above and infected with dsRED-*Lp*. After 2 hours of infection, cells were washed 3 times with KHM buffer (110 mM potassium acetate, 20 mM HEPES, and 2mM MgCl_2_, pH 7.3) and incubated for 1 min in KHM buffer with 50 μg/mL digitonin (Sigma-Aldrich). Cells were washed 3 times with KHM buffer and stained for 15 min at 37°C with primary antibody to *L. pneumophila* (1:1000 dilution; gift from Craig Roy) in KHM buffer with 3% BSA. Cells were washed with PBS, fixed, and quenched with 0.1 M glycine for 10 min. Cells were washed and incubated with secondary antibody anti-rabbit Alexa Fluor 488 for 1 hour, followed by washes and mounted on glass slides with DAPI mounting medium. Cells were analyzed by microscopy. 0.1% saponin in KHM buffer was used as a positive control for this assay. The percentage of infected cells harboring cytosolic bacteria out of the total number of infected cells were quantified.

### Statistical Analysis

GraphPad Prism software was used for graphing of data and all statistical analyses. Statistical significance for experiments with THP-1 cells was determined using the unpaired two-way Student’s *t* test. Statistical significance for hMDMs was determined using the paired two-way *t* test in experiments comparing multiple donors and the unpaired two-way *t* test in experiments involving infections with dsRED-expressing *Lp* for immunofluorescence assay. In hMDM experiments that compare cells from multiple donors, data are graphed so that each data point represents the mean of triplicate wells for each donor, and all statistical analysis was conducted comparing the means of each experiment. Differences were considered statistically significant if the *P* value was <0.05.

## Acknowledgements

We thank Shin lab members Natasha Lopes Fischer and Nawar Naseer as well as Igor Brodsky for critical reading of the manuscript. We also thank members of the Shin and Brodsky labs for helpful discussion and feedback. We thank Elisabet Bjanes for protocols and instruction on confocal microscope usage and imaging software analysis. We thank Gordon Ruthel for providing helpful training and advice on confocal microscopy. We thank the Human Immunology Core of the Penn Center for AIDS Research and the Abramson Cancer Center for providing purified primary human monocytes. We also thank the Penn Vet Imaging Core for allowing access and usage of the Leica SP5-II Confocal/Fluorescence Lifetime Imaging Microscope. This work is supported, in part, by NIH grant S10 RR027128-01 (to Penn Vet Imaging Core), NIH National Institute of Allergy and Infectious Diseases Grant R01AI121148 (to S.S.), National Science Foundation Graduate Fellowship DGE-1845298 (to A.R.B.), and a Burroughs–Wellcome Fund Investigators in the Pathogenesis of Infectious Diseases Award (to S.S.).

