## Supplemental Materials for "Human GBP1 promotes pathogen vacuole rupture and inflammasome activation during *Legionella pneumophila* infection"

**Figure S1**

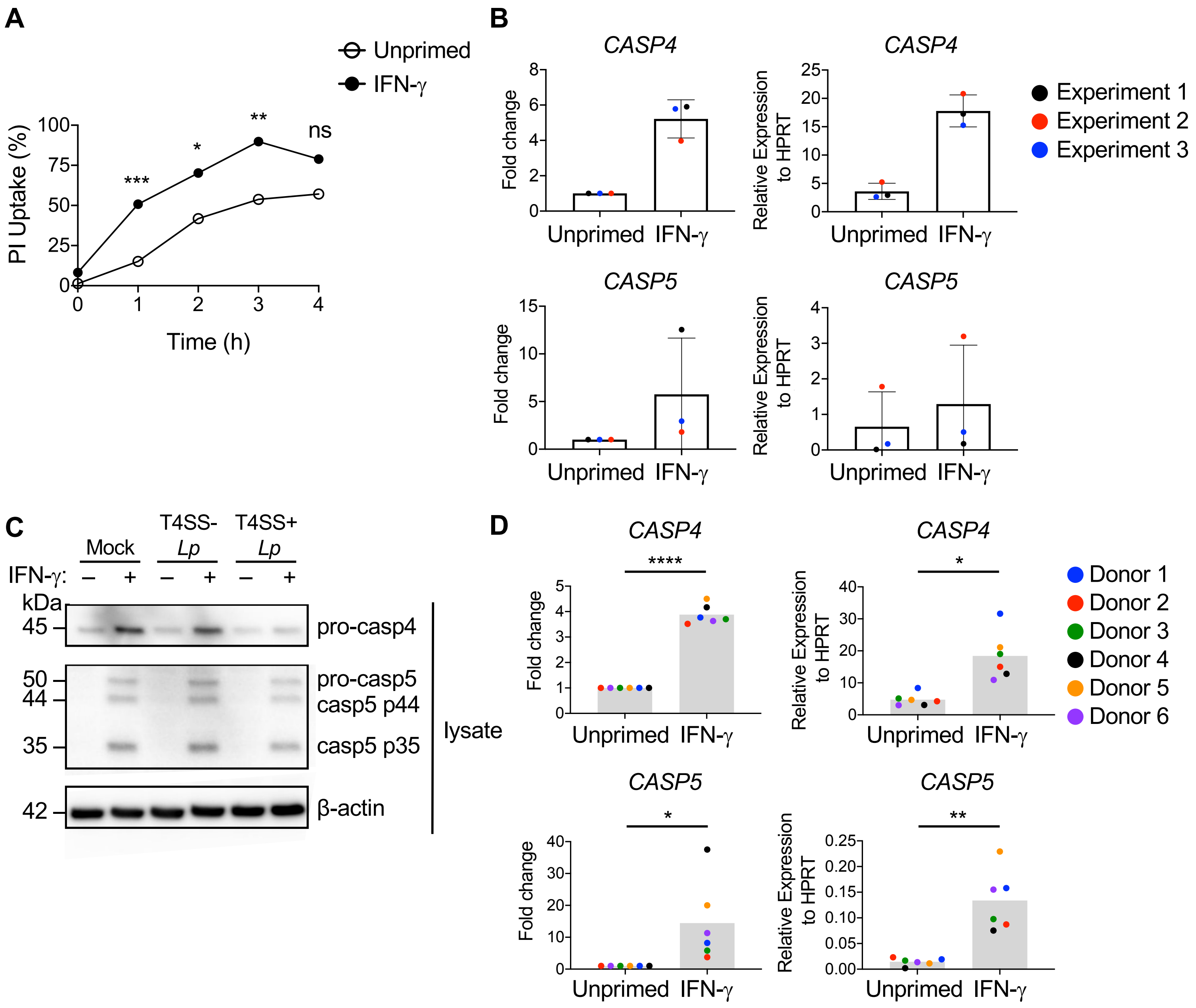

A
Unprimed
IFN-γ
PI Uptake (%)
Time (h)
***
*
**
ns
B
CASP4
CASP5
Fold change
Relative Expression to HPRT
Unprimed
IFN-γ
Experiment 1
Experiment 2
Experiment 3
C
Mock
T4SS- Lp
T4SS+ Lp
IFN-γ:
kDa
45
50
44
35
42
pro-casp4
pro-casp5
casp5 p44
casp5 p35
β-actin
lysate
D
CASP4
CASP5
****
*
**
Donor 1
Donor 2
Donor 3
Donor 4
Donor 5
Donor 6

#### Figure S2

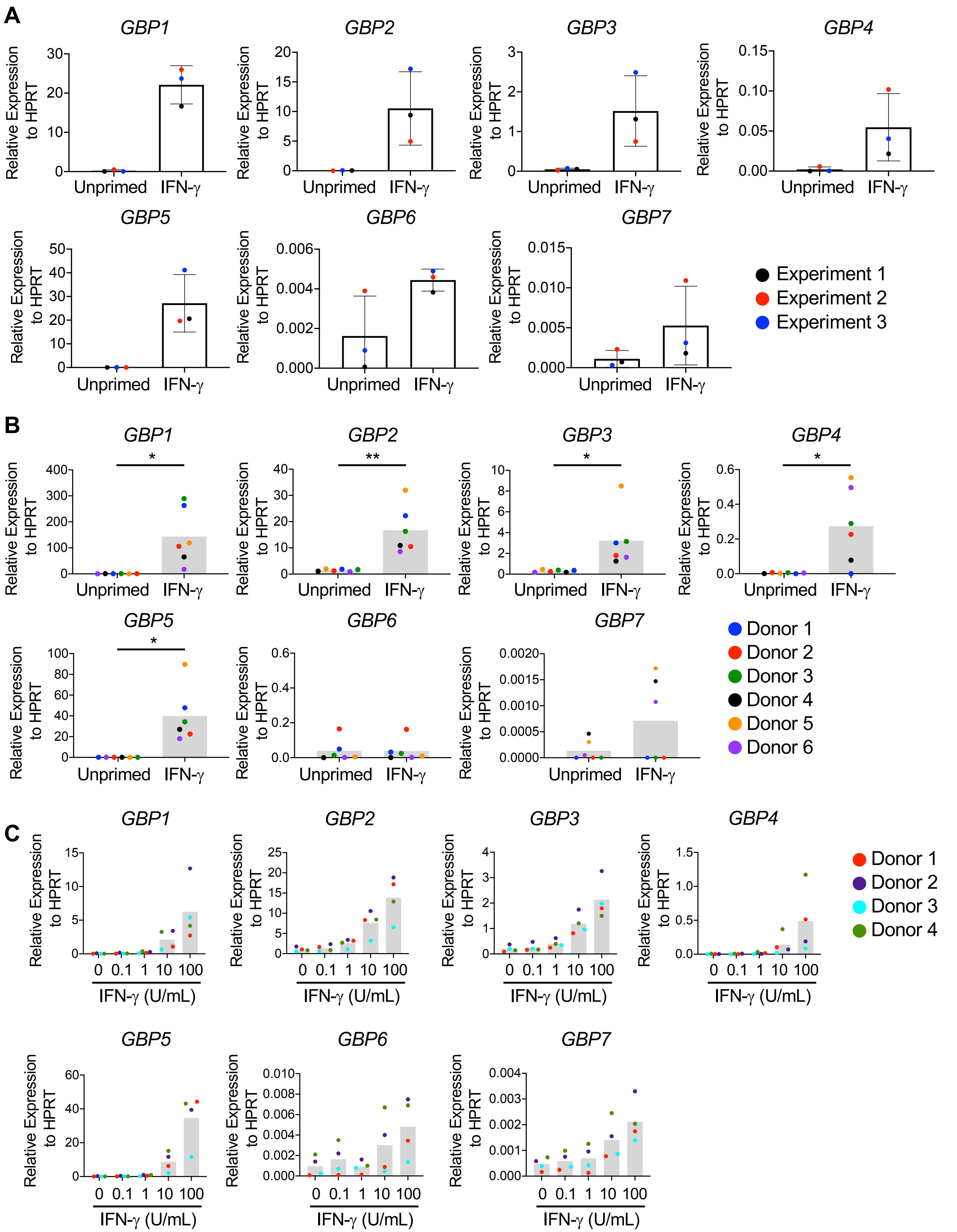

A
GBP1
GBP2
GBP3
GBP4
GBP5
GBP6
GBP7
Relative Expression to HPRT
Unprimed
IFN-γ
Experiment 1
Experiment 2
Experiment 3
B
Donor 1
Donor 2
Donor 3
Donor 4
Donor 5
Donor 6
C
IFN-γ (U/mL)
0
0.1
1
10
100

**Figure S3**

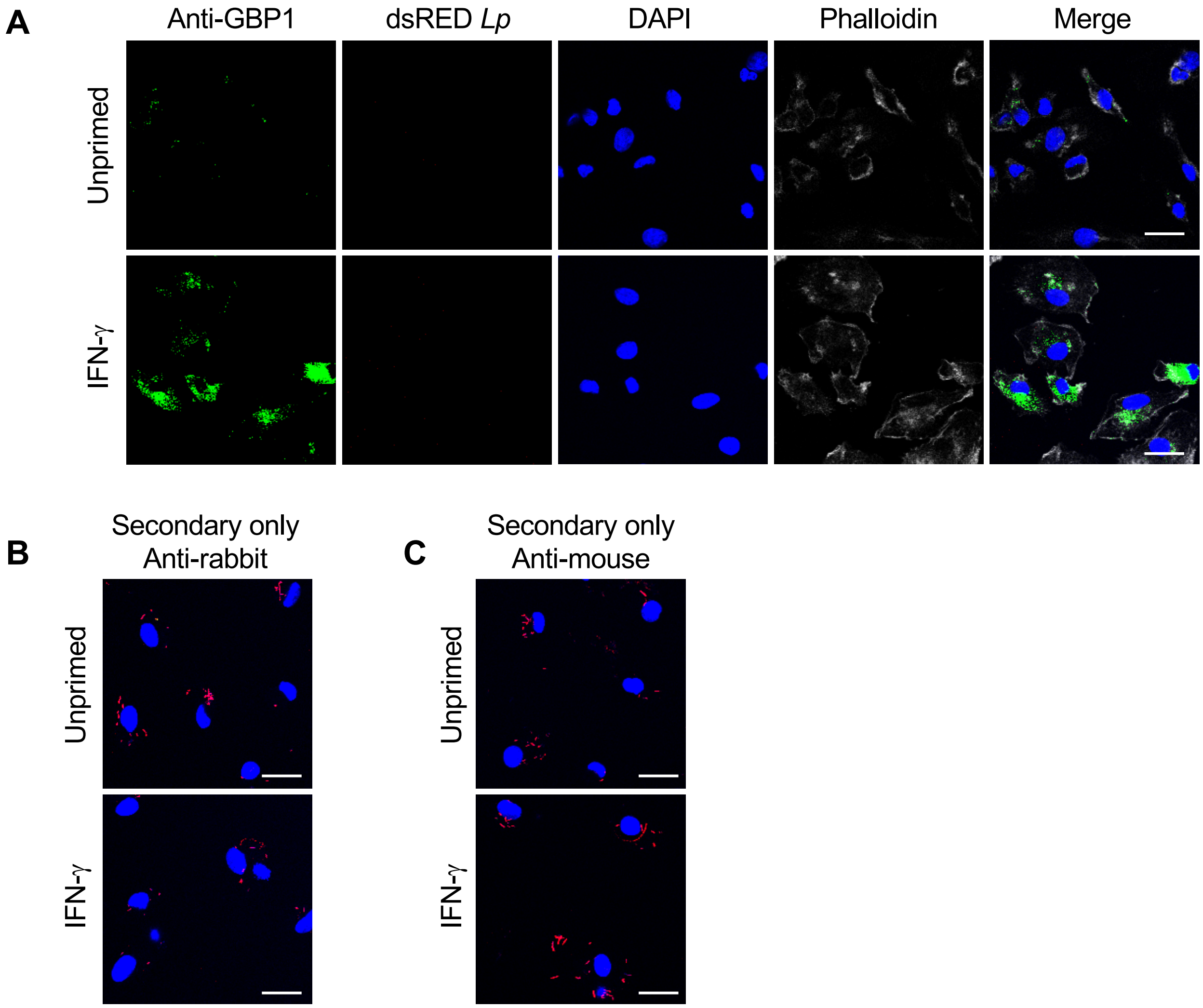
A
Anti-GBP1
dsRED Lp
DAPI
Phalloidin
Merge
Unprimed
IFN-γ
B
Secondary only
Anti-rabbit
Unprimed
IFN-γ
C
Secondary only
Anti-mouse
Unprimed
IFN-γ

### Figure S4

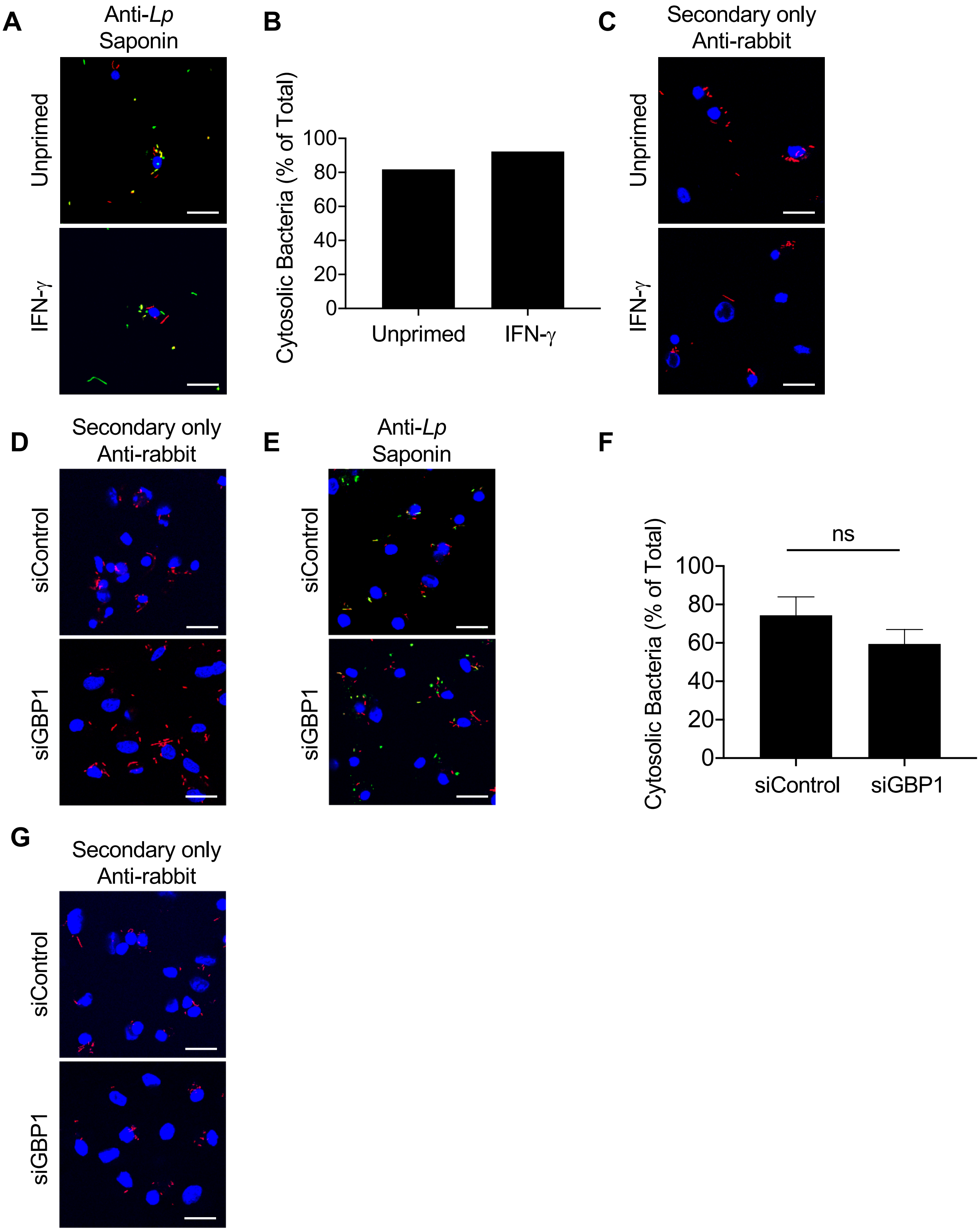

A
Anti-Lp
Saponin
Unprimed
IFN-γ
B
Cytosolic Bacteria (% of Total)
100
80
60
40
20
0
Unprimed
IFN-γ
C
Secondary only
Anti-rabbit
Unprimed
IFN-γ
D
Secondary only
Anti-rabbit
siControl
siGBP1
E
Anti-Lp
Saponin
siControl
siGBP1
F
ns
Cytosolic Bacteria (% of Total)
100
80
60
40
20
0
siControl
siGBP1
G
Secondary only
Anti-rabbit
siControl
siGBP1

**Supplementary Table 1:** qRT-PCR primers

| | |
|---|---|
| *HPRT1* forward | CCTGGCGTCGTGATTAGTGAT |
| *HPRT1* reverse | AGACGTTCAGTCCTGTCCATAA |
| *GBP1* forward | AGGAGTTCCTTCAAAGATGTGGA |
| *GBP1* reverse | GCAACTGGACCCTGTCGTT |
| *GBP2* forward | CTATCTGCAATTACGCAGCCT |
| *GBP2* reverse | TGTTCTGGCTTCTTGGGATGA |
| *GBP3* forward | ATTCCCTGAAGCTAACGCAAG |
| *GBP3* reverse | GGGCAGATCGAAGACAAAACATT |
| *GBP4* forward | ATGGGTGAGAGAACTCTTCACG |
| *GBP4* reverse | TGCGGTATAGCCCTACAATGG |
| *GBP5* forward | CCATGTGCCTCATCGAGAACT |
| *GBP5* reverse | ACAGGTTGCGTAATGGCAGAC |
| *GBP6* forward | ATGGAATCTGGACCCAAAATGTT |
| *GBP6* reverse | GCTGGTTCACCAATAGCTGCT |
| *GBP7* forward | TGCCTTCTTACCAAGTCCAGA |
| *GBP7* reverse | TCTCTGATGCCATGTTCAGG |
| *CASP4* forward | TCTGCGGAACTGTGCATGATG |
| *CASP4* reverse | TGTGTGATGAAGATAGAGCCCAT |
| *CASP5* forward | TCACCTGCCTGCAAGGAATG |
| *CASP5* reverse | TCTTTTCGTCAACCACAGTGTAG |

**Supplemental material**

**Fig. S1.** IFN-γ promotes inflammasome activation in response to *L. pneumophila* in human macrophages and upregulates caspase-4 and caspase-5. (A) PI uptake time course of PMA-differentiated THP-1 cells that were either left unprimed or primed with IFN-γ (100 U/ml) for 24 h and infected with T4SS+ *Lp* MOI=50*.* Data are representative of three independent experiments with each data point representing the mean of triplicate infected wells. *P< 0.05, **P< 0.01, ***P< 0.001 by unpaired t-test. PMA-differentiated THP-1 cells (B) or primary hMDMs (D) were either left unprimed or primed with IFN-γ (100 U/mL) for 18 or 20 hours, respectively. Transcript levels of *caspase-4* (*CASP4*) and *caspase-5* (*CASP5*) were determined by quantitative RT-PCR. Fold change was calculated by normalizing to the housekeeping gene HPRT for each sample and then to the unprimed sample. Relative expression of each gene was calculated by normalizing to the housekeeping gene HPRT. Shown are pooled results of three independent experiments (B) or six independent experiments using hMDMs from different healthy human donors (D), with each data point representing the value for each experiment (B) or an individual donor (D). *P< 0.05, **P< 0.01, and ****P< 0.0001 by paired t-test. (C) PMA-differentiated THP-1 cells were either left unprimed or primed with IFN-γ (100 U/ml) overnight and infected with T4SS- *Lp*, T4SS+ *Lp*, or mock-infected with PBS for two hours. Immunoblot analysis was performed on PMA-differentiated THP-1 lysates for full-length caspase-4 (pro-casp4), full-length caspase-5 (pro-casp5), caspase-5 intermediates (casp5 p44 and casp5 p35), and β-actin (same β-actin blot as shown in Fig. 1C since from same experiment). Western blots are representative of three independent experiments.

**Fig. S2.** Human GBPs are transcriptionally upregulated by IFN-γ in macrophages. PMA-differentiated THP-1 cells (A) or primary hMDMs (B) were either left unprimed or primed with IFN-γ (100 u/mL) for 18 or 20 hours, respectively. (C) hMDMs were left unprimed or primed with IFN-γ at the indicated concentrations for 20 hours. (A, B, C) Transcript levels of *GBP1-7* were determined by quantitative RT-PCR and relative expression of each gene was calculated by normalizing to the housekeeping gene HPRT. Shown are the pooled results of three independent experiments (A) or six independent experiments using hMDMs from different healthy human donors (B), with each data point representing the value for each experiment (A) or an individual donor (B). *P< 0.05 and **P< 0.01 by paired t-test. (C) Shown are the pooled results of four independent experiments using hMDMs from different healthy human donors and each data point represents the value of an individual donor.

**Fig. S3.** GBP1 is distributed throughout the cytoplasm in uninfected hMDMs. Primary hMDMs were either left unprimed or primed with IFN-γ (100 U/mL) overnight and infected with dsRED-expressing T4SS+ *Lp* or left uninfected for two hours. (A) Representative fluorescence micrographs of anti-GBP1 staining in uninfected hMDMs. (B and C) Representative fluorescence micrographs of dsRED-T4SS+ *Lp*-infected hMDMs stained with only secondary-antibody anti-rabbit (B) or anti-mouse (C) Alexa Fluor 488. (A, B, C) Images are representative of three independent experiments using hMDMs from different healthy human donors.

**Fig. S4.** Controls for phagosome integrity assay and GBP1 immunostaining assay. (A) Representative fluorescence micrographs of anti-*Lp* primary antibody and Alexa Fluor 488-conjugated anti-rabbit secondary antibody staining in saponin-permeabilized unprimed and IFN-γ-primed dsRED-T4SS+ *Lp*-infected hMDMs and (B) quantification of cytosolic *Lp*-infected hMDMs out of total infected hMDMs. (C) Representative fluorescence micrographs of unprimed and IFN-γ-primed dsRED-T4SS+ *Lp*-infected hMDMs stained with only secondary-antibody anti-rabbit Alexa Fluor 488 as a control for digitonin phagosome integrity assay. (D) Representative fluorescence micrographs of IFN-γ-primed siControl and siGBP1 dsRED-T4SS+ *Lp*-infected hMDMs stained with only Alexa Fluor 488-conjugated anti-rabbit secondary antibody as a control for GBP1 immunostaining assay. (E) Representative fluorescence micrographs of anti-*Lp* primary antibody and Alexa Fluor 488-conjugated anti-rabbit secondary antibody staining in saponin-permeabilized IFN-γ-primed siControl and siGBP1 dsRED-T4SS+ *Lp*-infected hMDMs and (F) quantification of cytosolic *Lp*-infected hMDMs out of total infected hMDMs. (G) Representative fluorescence micrographs of IFN-γ-primed siControl and siGBP1 dsRED-T4SS+ *Lp*-infected hMDMs stained with only Alexa Fluor 488-conjugated anti-rabbit secondary antibody as a control for digitonin phagosome integrity assay. (A-G) Data and images are representative of three independent experiments using hMDMs from different healthy human donors.
